# Hippocampal Asymmetry of Regional Development and Structural Covariance in Preterm Neonates

**DOI:** 10.1101/2020.11.25.397521

**Authors:** Xinting Ge, Yuchuan Qiao, Shiyu Yuan, Wenjuan Jiang, Mengting Liu

## Abstract

Premature birth is associated with high prevalence of neurodevelopmental impairments in surviving infants. The hippocampus is known to be critical for learning and memory, the putative role of hippocampus dysfunction remains poorly understood in preterm neonates. Particularly, hemispherical asymmetry of the hippocampus has been well-noted, either structurally or functionally. How the preterm birth impairs the hippocampal development, and to what extent the hippocampus was impaired by preterm birth asymmetrical has not been well studied. In this study, we compared regional and local hippocampal development in term born neonates (n=361) and prematurely born infants at term-born equivalent age on MRI studies (n = 53) using T2 MRI images collected from the Developing Human Connectome Project (dHCP); We compared 1) volumetric growth; 2) shape development in the hippocampal hemispheres using Laplace–Beltrami eigen-projection and boundary deformation between the two groups; and 3) structural covariance between hippocampal vertices and the cortical thickness in cerebral cortex regions. We demonstrated that premature infants have smaller volume for the right hippocampi, while no difference was observed for the left hippocampi. Lower thickness was observed in the hippocampal head in both hemispheres for preterm neonates compared to full-term peers, while an accelerated hippocampal thickness growth rate was found in left hippocampus only. Structural covariance analysis demonstrated that in premature infants, the structural covariance between hippocampi and limbic lobe were severely impaired compared to healthy term neonates only in left hemisphere. These data suggest that the development of the hippocampus during the third trimester may be altered following early extrauterine exposure, with high degree of asymmetry. These findings suggested that the hippocampus shows high degree of vulnerability, particularly asymmetrical vulnerability or plasticity, in preterm neonates at the term-born equivalent age compared to full-term healthy controls.

## Introduction

Across multiple neuroimaging investigations, alterations of the developmental trajectory in both global and region-specific brain structures have been demonstrated in premature infants compared to full-term (FT) peers (Smitthimedhin et al., 2018; Wu et al., 2017). Hippocampus is the central node in mnemonic circuitry, and yield a dramatic morphological alteration in the 3^rd^ trimester, which is also the period when the preterm births occur (Kim et al., 2020). Preterm birth has demonstrated impaired growth (Beauchamp et al., 2008; Thompson et al., 2008) and distinct folding patterns of the hippocampus (Thompson et al., 2013). Such changes may be associate with a wide range of adverse developmental outcomes including memory deficits compared with term-born controls, which could severely impact individual’s daily life in the future (Nosarti et al., 2016; Nosarti et al., 2014; Thompson et al., 2020). Investigating the neuroanatomical development trajectory of preterm hippocampi at term-born equivalent time period, could potentially contribute to the prediction and early intervention of the adverse outcomes.

Morphometry study based on volume analyses has been performed to reveal the development course of the hippocampus during the perinatal period. Linear increase of hippocampal volume was found in healthy fetuses (Jacob et al., 2011). For preterm neonates, the hippocampal volume has been shown to be more symmetrical compared to full-term infants (Thompson et al., 2008). However, the development of hippocampus is not unitary but rather heterogeneous (Tanaka, 2020). Hemispherical asymmetry of the hippocampus has been well-noted (Bajic et al., 2012; Ge et al., 2015; Thompson et al., 2009), either structurally or functionally (Krogsrud et al., 2014; Sakaguchi and Sakurai, 2017; Uematsu et al., 2012). Left–right asymmetries have likely evolved to make optimal use of bilaterian nervous systems (Shipton et al., 2014). Recent work found that the structural and functional divergence between hippocampal hemispheres are supported by synaptic size and shape, neuronal circuit connections and mechanisms according to brain hemisphere (Shinohara et al., 2008; Shipton et al., 2014). Hence, if preterm birth exerts influence on hippocampal development, we might expect more heterogeneously and asymmetrically morphological alterations due to the nature of neuronal architecture of the hippocampus.

Shape analyses method, on the other hand, has the potential to identify local changes or developmental states of brain (sub)regions, which is insufficient by using volume analysis (Gerig et al., 2001). The tendency of hippocampal symmetry in preterm neonates (Thompson et al., 2008) compared to full term neonates might be associate with its global or sub-regional vulnerability to a wide variety of neurological insults related to preterm birth, including hypoxic-ischemic injury, stress (Perlman, 2001) and metabolic disturbance (Schmidt-Kastner and Freund, 1991). Shape analysis may yield local information related to the neurological insults during the development of the hippocampus which is unavailable previously (Gerig et al., 2001; Ho and Magnotta, 2010; Joseph et al., 2014; Wright et al., 1995).

In addition, the structural covariance (SC) approach provides an effective way to characterize inter-regional associations of morphological properties (DuPre and Spreng, 2017; Khundrakpam et al., 2017; Li et al., 2013; Mechelli et al., 2005). The hippocampus is not a homogeneous structure, with both structural and functional differences along its longitudinal axis (Poppenk et al., 2013). Posterior (pHC) and anterior (aHC) hippocampus has been extensively reported to have differently structural covariance to different cerebral lobes, which were affected by preterm birth (Ball et al., 2012; Kim et al., 2020). However, it remains unknown to what extent the structural covariance between hippocampus and cerebral cortex can be affected. With the help of high-resolution MRI and shape analysis method, it is feasible to conduct vertex-wise structural covariance to cortical areas, which may provide more detailed developmental hippocampal trajectories, either alone or association with cortical regions.

Past studies indicated that different subsections of the hippocampus follow different developmental trajectories. To what extent preterm birth could influence the developmental trajectories, for both left and right hippocampus, is the main target of this studies. Specifically, The aims of this study were to determine: 1) differences in hippocampal volumes asymmetry between full term and preterm infants at term equivalent age; 2) differences in hippocampal shapes asymmetry between full term and preterm neonates at term equivalent age, especially the shape development; and 3) associations between hippocampal shape, specifically thickness, and cortical thickness and their differences shape. The hypotheses were that: 1) preterm neonate hippocampi would have altered volume and shape compared with full term neonate hippocampi, with changes relating regionally altered hippocampal thickness; 2) alterations to preterm hippocampal volumes would be related to developmental trajectory in regional shapes; and 3) developmental trajectory alterations in preterm birth neonates may reflect the structural covariance map for hippocampal thickness.

## Materials and Methods

### Participants

Hippocampus were extracted from T2-weighted MRI scans of subjects participated in the developing Human Connectome Project (dHCP). Written informed parental consent was obtained from all participants before enrolment to the study. Datasets acquired from only singletons were included in this study (86 twin images were excluded). Preterm neonates scanned before 37 weeks (n = 49) were also excluded. The rest images were visually inspected and datasets with substantial motion on MRI or major focal parenchymal lesions at the time of their first scan were excluded. The final sample consisted of 414 (163 female) neonates ranging in age at scan (post-menstrual age; PMA) from 37 to 44 weeks, with a gestational age (GA) at birth of 24–42 weeks (term: mean PMA 40.0 weeks, mean GA 41.1 weeks; preterm: mean PMA 32.1 weeks, mean GA 40.2 weeks).

### MRI acquisition

All datasets were acquired on a Philips Achieva 3.0 T scanner at the Evelina Newborn Imaging Centre using a dedicated 32-channel neonatal head coil (Batalle et al., 2017). All anatomical volumes were collected as part of the dHCP and are described in detail in (Makropoulos et al., 2018).

For T2-weighted anatomical scans, turbo spin echo (TSE) sequences were used with two stacks per weighting, sagittal and axial. For T2-weighted scans, the parameters were: repetition time = 12 s, echo time = 156, SENSE = 2.11 (axial) and SENSE = 2.58 (sagittal). Images were acquired with a repetition time = 4.8 s, echo time = 8.7, inversion time = 1740, SENSE factor of 2.26 (axial) and 2.66 (sagittal). For all images the in-plane resolution was 0.8 × 0.8 mm with a slice thickness of 1.6 mm, with a slice overlap of 0.8 mm. The resulting images were motion corrected as described in (Cordero Grande et al., 2018) and super-resolution reconstruction was performed as in (Kuklisova-Murgasova et al., 2012), resulting in 3D volumes resampled to 0.5 mm isotropic resolution. The resulting images were also corrected for bias field inhomogeneities.

### Image preprocessing

Intensity inhomogeneity correction was firstly applied to all the images using the N3 algorithm. Removal of the non-brain tissue for each individual was performed using a deep learning-based brain segmentation approach, i.e. 3-D Unet (Hwang et al., 2019) based on 12templates (T2 MRI + manually segmented brain masks) in dHCP dataset. The 3D-Unet consists of three contracting encoder layers to analyze the whole image and three successive expanding decoder layers to produce a full-resolution segmentation (Ronneberger et al., 2015). Each layer contains two 3 × 3 × 3 convolutions each followed by a rectified linear unit (ReLu), and then a 2 × 2 × 2 max pooling with strides of two in each dimension. The 3D-Unet was trained on 36 × 36 × 36 voxels image patches fragmented from template MRI images. After the brain extraction, a whole brain template was then calculated using Advanced Normalization Tools (ANTs) (Avants et al., 2009). The script *buildtemplateparallel.sh* (Avants and Gee, 2004) was run for the whole cohort using the default setting in ANTs.

### Segmentation of the hippocampus

Manual segmentation of the hippocampus was conducted on the brain template using ITK-SNAP software (Yushkevich et al., 2016), reference by the segmentation protocol of the fetal hippocampus in our previous publication (Ge et al., 2015). The masks of both the left and right hippocampi generated in the template space were then reversely registered to the individual space using the deformation information during the template construction process. The individual masks of the hippocampi were then visually checked and modified manually by an expert anatomist.

### Shape Analysis

Shape analysis method has been used in our previous studies to assess the developmental trajectories of the fetal hippocampus and cerebellum (Ge et al., 2015). In the current research, the binary masks of each hippocampus including the template were firstly converted to triangular meshes. The spurious features caused by segmentation artifacts were detected and removed using iterated Laplace–Beltrami (LB) eigen-projection and boundary deformation (Shi et al., 2010). The resulting surface meshes were then remeshed to 1000 uniformly distributed vertices, which was determined according to the size of the hippocampus during the perinatal period as well as considering the calculation cost.

The individual triangulated meshes were then registered to the template mesh using the novel curvature-driven surface mapping algorithm, Riemannian metric optimization on surfaces (RMOS), which incorporated both geometric and anatomical features to guide surface mapping in the LB embedding space (Gahm et al., 2018). Detailed one-to-one correspondences across different surfaces of the hippocampi were established, which can achieve superior performance in the prediction of hippocampal subfield changes compared with our previous approach (surface mapping algorithm) via conformal metric optimization on surfaces (CMOS). Similar to the previous studies, the thickness measured at each vertex of the mapped surface was defined as the distance from the vertex to the medial core of the hippocampus. Left and right hippocampi were calculated separately.

### Cortical thickness of brain lobes

Cortical thickness of neonatal MRI is calculated using the NEOCIVET 2.0 pipeline (Liu et al., 2019); Specifically, the pipeline begins with general data pre-processing, including denoising and intensity nonuniformity correction. After the brain extraction using 3D-Unet (Hwang et al., 2019), the skull-striped brain was registered to the MNI-NIH neonatal brain template (http://www.bic.mni.mcgill.ca/ServicesAtlases/NIHPD-obj2), with spatial resolution of 0.6 × 0.6 × 0.6 mm^3^ (isotropic). Different types of brain tissue (GM, WM, and CSF) are thereafter segmented by an advanced deep non-local 3D-Unet (Wang et al., 2020). Individual templates (MRI + manually segmented tissue labels) utilized for the deep learning approach are then selected evenly across all postmenstrual ages (PMA). Next, the corpus callosum is segmented on the midline-plane and used to divide the WM into hemispheres. Then a marching-cube based framework is adopted to generate a triangulated mesh WM surface attached to the boundary between the GM and WM. After resampled to a fixed number of 81,920 surface meshes (triangles) using the icosahedron spherical fitting, this surface is further fitted to the sharp edge of the GM-WM interface based on the image intensity gradient, which preserves the spherical topology of the cortical mantle. A CSF skeleton is then generated from the union of WM and CSFs. Pial surface is constructed by expanding the WM surface towards the skeleton as an intermediate pial surface. The intermediate pial surface further undergoes a fine deformation to identify actual edges of sulcal CSF volumes using an intensity gradient feature model. Finally, the cortical thickness is estimated based on the Euclidean distance between the white matter and pial surface, with a smooth kernel size of 10 mm (Vasung et al., 2016).

### Structural covariance

A general linear model (GLM) was used to account for structural covariance between thickness on hippocampal vertices and vertices in cortical lobes while accounting for confounding variables including PMA at birth, PMA at scan, sex, and brain volume. Next, correlation matrices between cortical thickness in specific anatomical cortical lobe and hippocampus across the whole hippocampal vertices were computed (Lerch et al., 2006; Raznahan et al., 2011). Results yielded a 6 x 2000 correlation matrix representing the structural covariance between average cortical thickness in 6 cortical lobes and 2000 hippocampal vertices.

The comparison of structural covariance between full term and preterm neonates were performed using permutation tests with 5000 repetitions. Specifically, in each run of permutation, all neonates were randomly assigned into the two subgroups. The number of neonates within each group was same as two subgroups in real conditions. Structural covariance was then generated and compared from randomly assigned neonates. For each variable (lobes on cortical surface or vertices on hippocampal surface), the actual difference between the given two groups was placed in its corresponding permutation distribution to obtain the significance level (95%). Finally, the false discovery rate (FDR) of P-values from all variables and was controlled (Benjamini and Hochberg, 1995).

We further analyzed the structural covariance using a ‘winner take all’ strategy, where the cortical subdivision that correlates strongest with a given hippocampal voxel ‘wins,’ (Zhang et al., 2008). This strategy was widely used in thalamus connectivity studies in past literatures (Kim et al., 2013; Mills et al., 2012). This approach partitioned the hippocampus into distinct subdivisions. The spatial organization of these hippocampal subdivisions were compared between full term and preterm neonates to qualitatively evaluate the influence of early extrauterine exposure on the development trajectory of hippocampus.

## Results

### Volume growth of hippocampus

Using general linear model (GLM), including PMA at birth and PMA at scan as covariates, hippocampal volume was found larger in right hemisphere for both term (volume: p < 0.0001; normalized volume: p < 0.0001) and preterm (volume: p = 0.0052; normalized volume: p < 0.0001) groups.

Hippocampal volume and normalized hippocampal volumes in relation to the age are shown in Fig. 1. Each point denotes the hippocampal volume of one subject. Solid line denotes linear growth curve for volumes in one group. Overall, volumes (p = 0.041) and normalized (p = 0.0072) volumes of the right hippocampal hemispheres in premature infants were smaller compared to terms peers. Volume difference was not found in left hippocampi.

**Figure 1.**
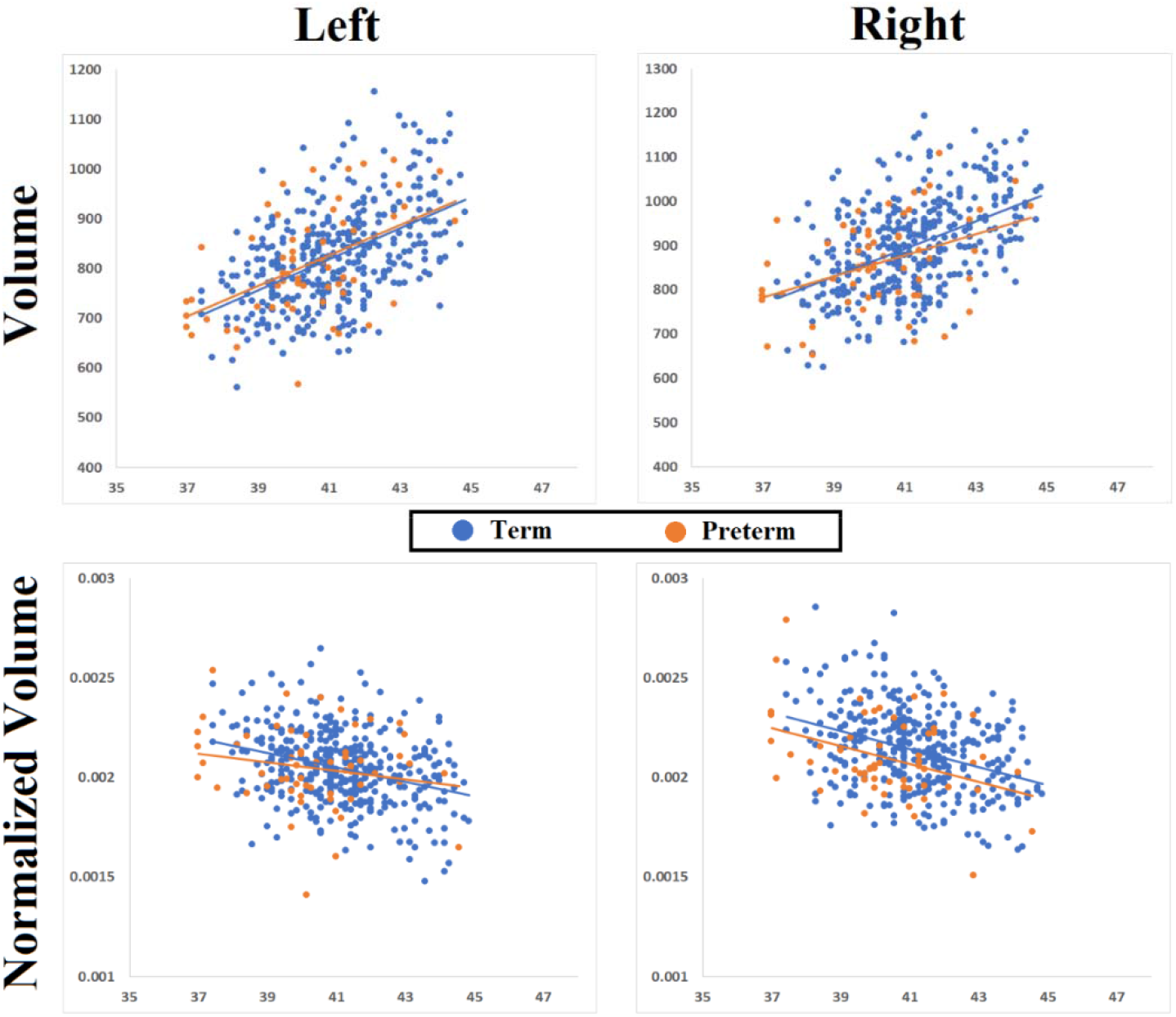
Volume of left and right hippocampus in term and preterm neonates

The growth rates (cm^3^/week) in both hemisphere of the hippocampus for both term and preterm infants from 30 to 46 weeks are shown in Table 1. The right hippocampus grows slower for preterm infants compared to term neonates. It is worth noting that after correction by the whole brain volume, volume of preterm hippocampus showed faster increase compared to term hippocampal volume.

**Table 1.**
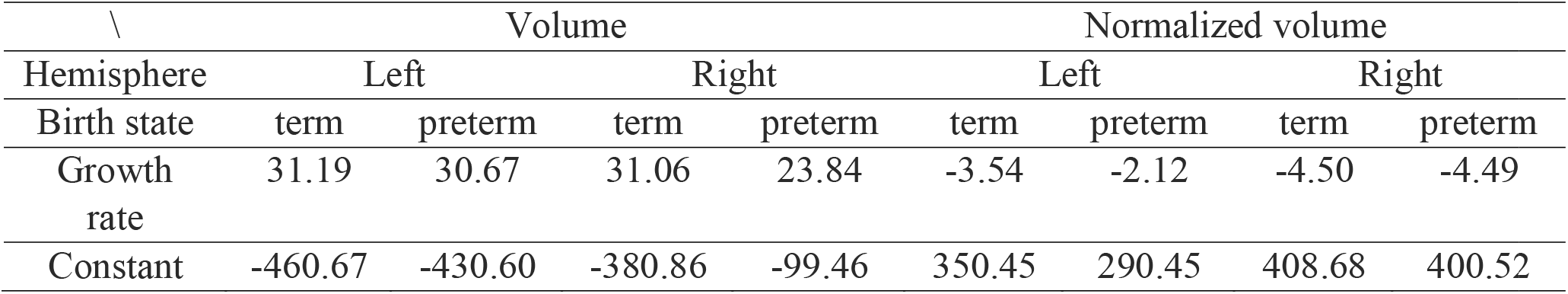
Comparison of hippocampal volume and volume growth rate between term and preterm neonates

### Shape difference

Fig. 2 shows four views (i.e., medial, lateral, superior, and inferior) of the hippocampal surface, with t-maps overlaid showing the comparison between the term and preterm groups. When controlling for GA, BA and brain volume at MRI, premature infants had significantly lower thickness in the medial part of hippocampal head for both hemispheres.

**Figure 2.**
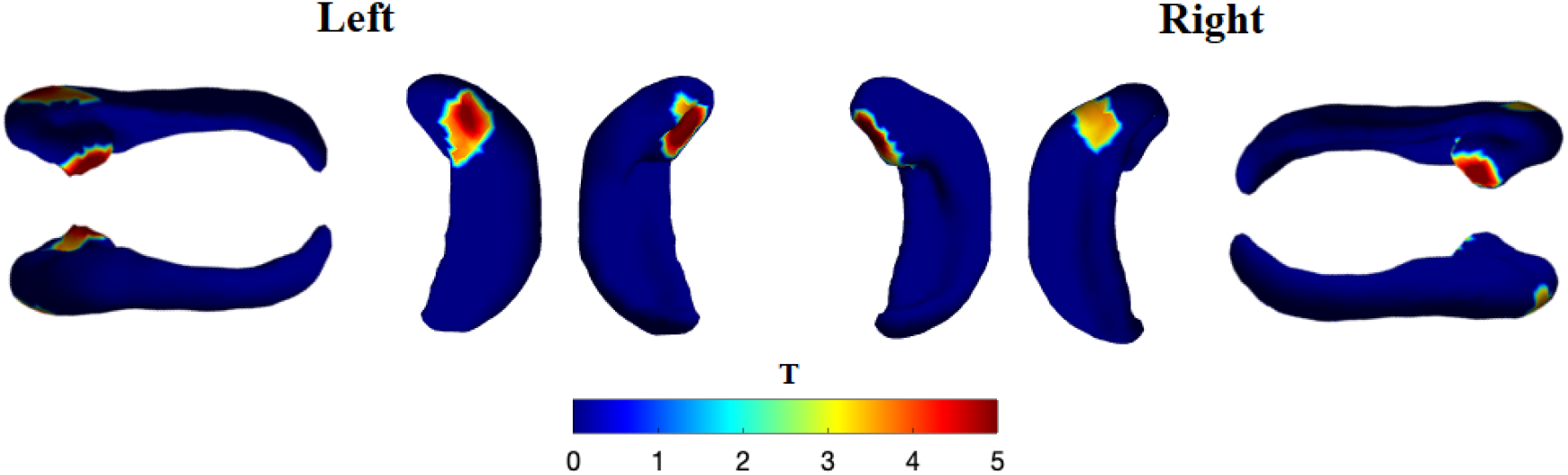
Thickness difference between term and preterm.

### Shape developmental trajectory

Images depicting shape development of the hippocampus in premature infants and healthy controls are shown in Fig.3. Different colors overlaid on the hippocampal surface indicate different growth rates, where red regions represent higher growth rates and blue regions indicate lower growth rates. The medial and lateral parts of the hippocampus on both left and right hemispheres showed more rapid expansion than the superior and inferior parts in both premature infants and controls. The interactive significance of regional thickness growth in terms of PMA (after controlling for GA, BA and brain volume, same below) is estimated using SurfStat toolbox and no significant difference in thickness growth rate were found between term and preterm hippocampus. The t score between preterm and term neonates in each of the vertices were shown in Fig. 3. It is shown that interior and exterior lateral hippocampal regions exhibited faster growth rate.

**Figure 3.**
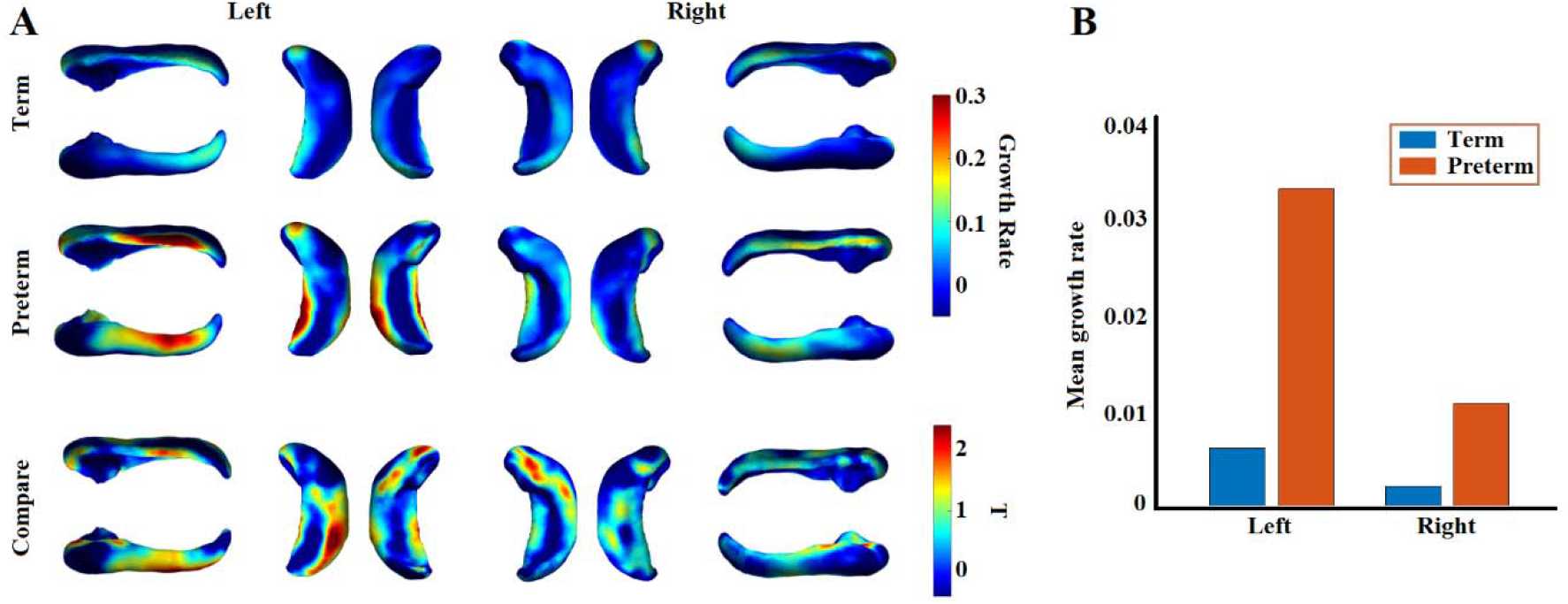
Growth rate of thickness in term and preterm neonates.

Although difference in growth rate was not found locally, globally, it was observed that averaged thickness across vertices on left hippocampal hemisphere showed faster growth rate in preterm neonates (0.033 mm/week) compared to term born controls (0.006 mm/week). Wheras, in right hippocampal hemisphere, preterm neonates (0.011 mm/week) didn’t show big difference compared to term born controls (0.002 mm/week). To statistically test whether the growth rate was larger in preterm neonates in left and right hippocampal hemisphere, a permutation tests with 5000 repetitions was conducted. In each repetition, all neonates were randomly assigned into the two subgroups with the same numbers as two subgroups in real conditions. Results revealed preterm neonates showed significantly higher growth rate in comparison to term born controls only in left hippocampal hemisphere (*p* = 0.011, 5000 permutation test), but not in right (*p* > 0.05, 5000 permutation test).

### Stronger hippocampal-limbic structural covariance was disrupted in preterm neonates only in left hemisphere

Average structural covariance between the hippocampus and 6 cortical lobes across all the vertices on hippocampal surface for left and right hemispheres respectively were plotted in Fig. 4. For term born infants, structural covariance between hippocampus and limbic cortex showed much higher values compared to that between hippocampus and other cortical lobes (p’s < 0.0001). The stronger SC between limbic cortex and hippocampus were found only in right hippocampal in preterm infants (p < 0.0001). We also compared the SC difference between term and preterm infants. Permutation test showed that on average, preterm neonates exhibited significantly less SC between hippocampus and limbic cortex (corrected p = 0.034) only for left hemisphere.

**Figure 4.**
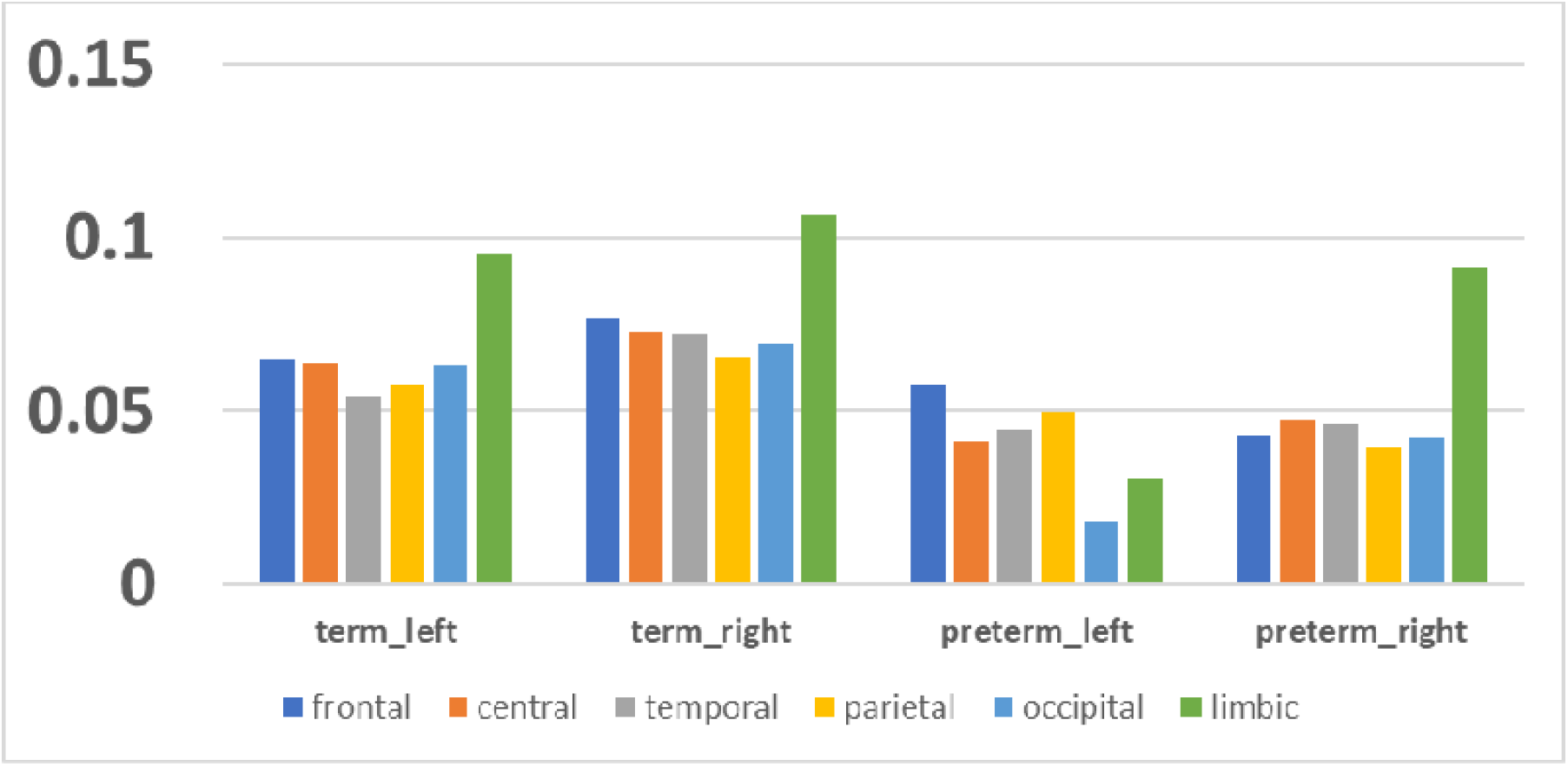
term-born neonates and preterm have higher structural covariance with limbic cortex.

**Figure 5.**
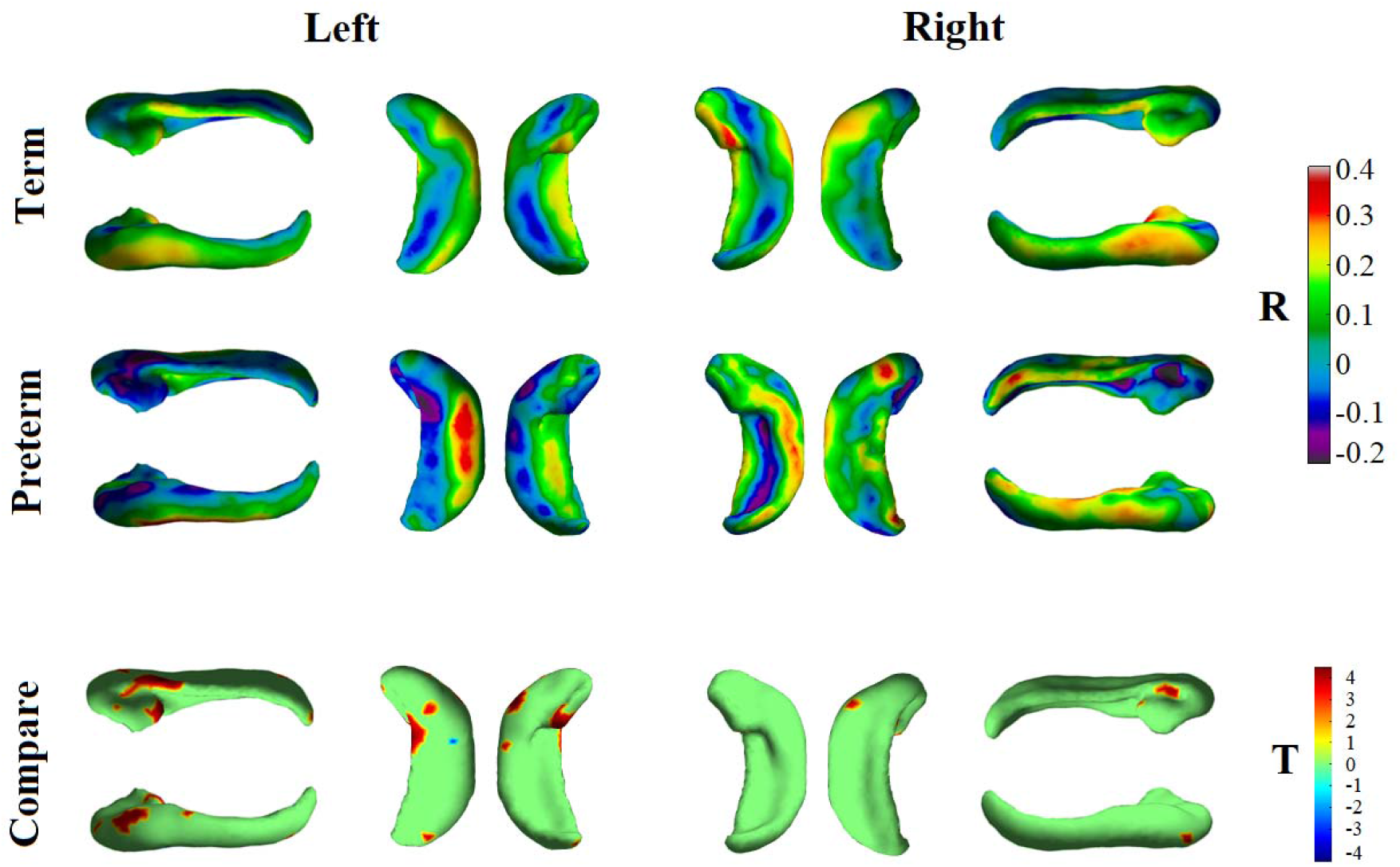
Hippocampal-limbic structural covariance comparison between full term and preterm neonates.

### Vertex-wised hippocampal-limbic structural covariance comparison between full term and preterm neonates

Images depicting hippocampus-limbic structural covariance values across hippocampal vertices in premature neonates and healthy controls are shown in Fig.5. The difference of the hippocampus-limbic SC evaluated using permutation test was shown in fig.5B. Red regions represent higher SC with limbic lobe cortical thickness in term controls, and blue regions indicate higher SC with limbic lobe cortical thickness in preterm infants. Significantly higher SC with limbic lobe was found mainly in vertices on anterior and posterior hippocampal head. Particularly, left hippocampal hemisphere exhibited more vertices with significantly higher SCs then right hippocampal hemisphere.

### Winner take all test

Multiple hippocampal SC maps were condensed into a single winner-take-all projection (Fig. 6) in which every hippocampal vertex is labeled with the color of the cortical lobes that generated the highest structural covariance value in thickness. Results exhibited that for left hippocampal hemisphere, 60.1% of the vertices were most strongly covariate with cortical thickness in limbic lobe for term born infants; Whereas for preterm neonates, only 27.4%. In right hippocampal hemisphere, 55.6% of the vertices for term and 42.7% of the vertices for preterm infants most strongly correlated with limbic cortex. This means that in left hippocampal hemisphere, large number of vertices yielded much impaired structural connectivity with limbic cortex in preterm infants compared to full term controls. However, this doesn’t occur, at least less severely occurs, in right hemisphere. Results suggested the SC between limbic lobe and hippocampus was asymmetry impaired by preterm birth, where left hemisphere was more disrupted to limbic cortex compared to right hemisphere. Specifically, the some of the disrupted regions were localized in superior and inferior hippocampal fissure regions in left hippocampus, which is exactly the regions showed faster growth rate in thickness compared to controls (see Fig. 2).

**Figure 6.**
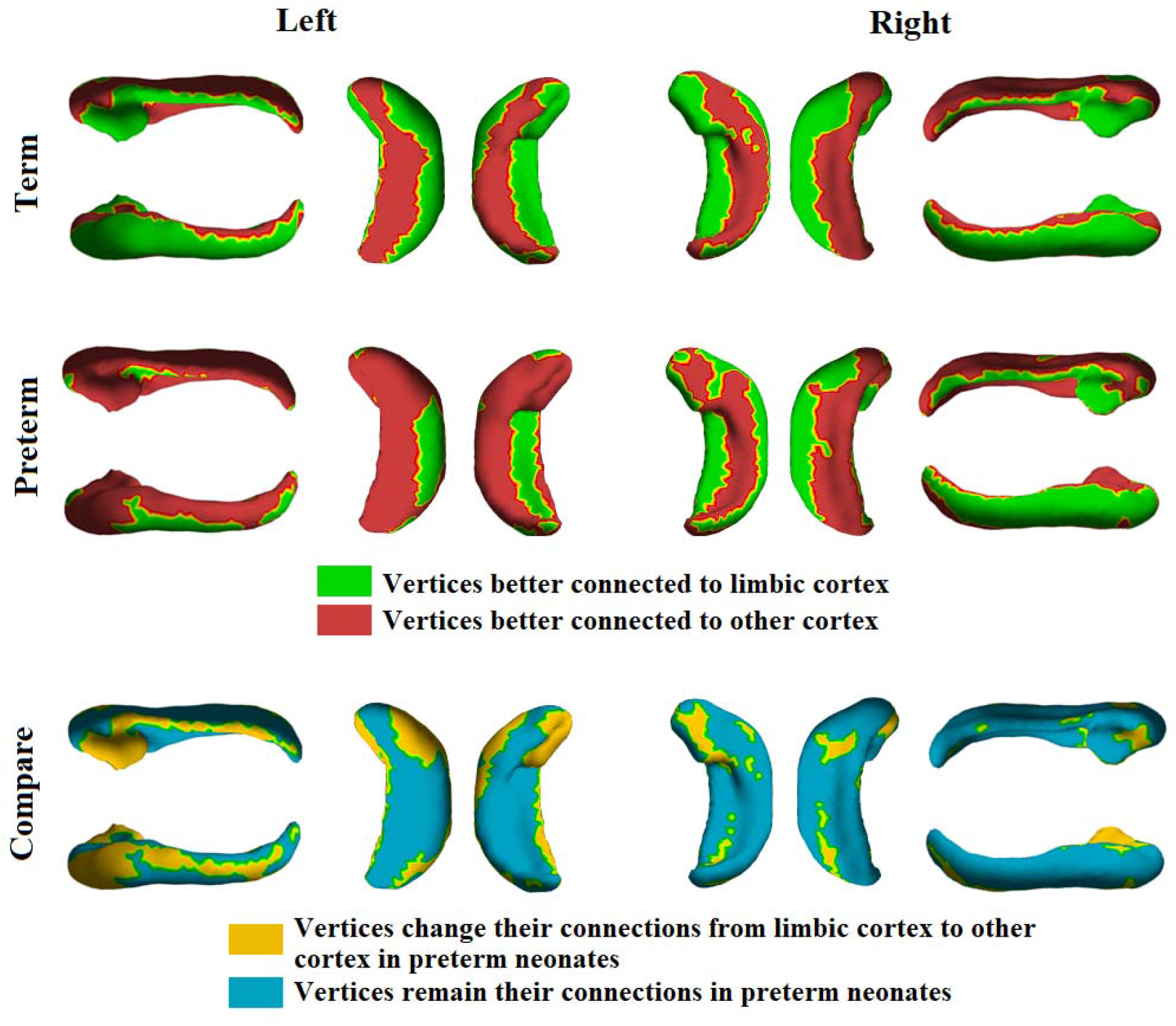
More vertices in term neonates have stronger structural covariance with limbic cortex. For preterm neonates, vertices that switched their connection from limbic cortex to other cortex were mostly found in left hippocampi.

### cortical thickness in 6 cortical regions doesn’t have difference

Unusual hippocampi-limbic structural covariance in preterm neonates may result from impaired limbic cortex development trajectory or hippocampal development trajectory, or both of them. Before we provided evidence that hippocampal development was damaged by preterm birth. To verify whether preterm birth also corrupt the cortical development in limbic cortex, we further compared the cortical thickness in limbic cortex between term and preterm neonates. Specifically, comparison of cortical thickness, cortical thickness growth rate, and structural covariance between limbic cortex with all other cortices were all performed. Statistical analysis utilized was identical to what was applied in hippocampal analysis. That is, omparison of cortical thickness and cortical thickness growth rate were conducted by GLM, structural covariances were compared using permutation tests with 5000 repetitions. By excluding the effects of birth age, scan age and brain volume, results indicated that none of the metrics were significantly different between the two groups, indicating that the preterm birth has very limited influence on the developmental trajectory of cortical thickness in limbic cortex in this period. The details of measures can be found in table 2.

**Table 2.** Comparison of cortical thickness in limbic cortex between term and preterm neonates

|  | CT | GR | SC-frontal | SC-central | SC-temporal | SC-parietal | SC-occipital |
| --- | --- | --- | --- | --- | --- | --- | --- |
| Term | 1.577 | 0.036 | 0.758 | 0.705 | 0.797 | 0.739 | 0.725 |
| Preterm | 1.581 | 0.041 | 0.709 | 0.735 | 0.746 | 0.740 | 0.748 |
| p-value | 0.872 | 0.414 | 0.788 | 0.813 | 0.576 | 0.801 | 0.769 |

## Discussion

In this study, using shape analysis and structural covariance methods, we found the hippocampus shows high degree of vulnerability, particularly asymmetrical vulnerability or plasticity, in preterm neonates at the term-born equivalent age compared to full-term healthy controls. We demonstrated that premature infants have smaller volume for the right hippocampi, while no difference was observed for the left hippocampi. Lower thickness was observed in the hippocampal head in both hemispheres for preterm neonates compared to full-term peers, while an accelerated hippocampal thickness growth rate was found in left hippocampus only. Structural covariance analysis demonstrated that the hippocampal thickness is significantly correlated with the cortical thickness of limbic lobe for both pre-term and full-term neonates compared to other cortical regions. However, in premature infants, the structural covariance between hippocampi and limbic lobe were severely impaired compared to healthy term neonates only in left hemisphere. These data suggest that the development of the hippocampus during the third trimester may be altered following early extrauterine exposure, with high degree of asymmetry.

### Asymmetrically altered hippocampal development trajectory by preterm birth

Hemispherical asymmetry of the hippocampus has been well-noted across the whole life span of human beings (PEDRAZA et al., 2004). The larger volume of right hippocampus was commonly found beginning *in utero* (Bajic et al., 2012) as early as from the early second trimester (Ge et al., 2015) to elderly aging populations (Wolf et al., 2001). Similar finding (larger right hippocampal volume) was also seen in current study for neonates in term-born equivalent age, regardless of the preterm birth.

Apart from the left-right hippocampal volume difference, we showed that preterm birth may alter hippocampal development asymmetrically in several unique and novel manners, which is seldom reported in past hippocampal asymmetry studies. These findings in turn may help outline the mechanism of how the preterm birth exert influence on the hippocampal development. First, smaller volume was found in premature neonates compared to term controls in right hemisphere, while no difference was found in left hippocampi. This may be interpreted by two possible schemes. One is right hippocampi is more vulnerable to the preterm birth induced impairments and is hence more easily contaminated by preterm birth. An alternative possible scheme is hippocampal development in both hemispheres are equally impaired by preterm birth, but only the left hippocampi express an “catch up” process, which drives a faster post-natal growth, leading to the normative left hippocampal volume. Using shape analysis, we found that preterm hippocampal thickness represents an accelerated growth in left hippocampi only. This provides a direct support for the “left hippocampal catch up” hypothesis rather than the “right hippocampal impairment” hypothesis. Post-natal accelerated growth in preterm neonates has been widely reported, which was thought to facilitate the preterm neonates to approximate the intra-uterine fetal growth patterns in extra-uterine environment due to cumulative protein and energy deficits (Raaijmakers and Allegaert, 2016). A recent study found in third trimester, cortical sulcation in cerebral cortex for preterm neonates exhibit a significant catch up process (Kim et al., 2020). The faster growth in left hippocampus may indicate a compensation, brain plasticity, or recovery (Kim et al., 2020) process that facilitate the left hippocampus to catch up the normal developmental trajectory.

Other than hippocampal volume and shapes, structural covariance also showed asymmetrical patterns in preterm neonates, i.e. preterm hippocampus exhibited impaired structural covariance to limbic cortex in left hemisphere only. Due to their adjoining anatomical locations, hippocampi-limbic connectivity (limbic cortex comprises the bilateral temporal pole lobe, cingulate cortex and para-hippocampal gyrus) has been commonly found higher than connectivity between hippocampus and other cortical regions from multiple studies using different modalities, e.g. tractography based white matter connectivity (Maller et al., 2019), structural covariance studies in adult brain (Guo et al., 2020; Nordin et al., 2018; Persson et al., 2014), and functional network studies in resting state (Kahn et al., 2008). The hippocampal-limbic connection also overlaps well with the structural connections of the direct hippocampal pathway (Duvernoy, 2005). The hippocampus and limbic cortex, especially the cingulate cortex and parahippocampal gyrus were both involved in an extended mnemonic system, namely Papez circuit (Shah et al., 2012), which was thought work interactively to facilitate the neural decoding of memory, learning, and emotion skills (Aggleton et al., 2016; Vann, 2013). The impaired hippocampal-limbic connectivity may reflect the influence of preterm birth to much wider neuro-functional development, and such influence starts from very early stages and extend to the perinatal periods during the brain development.

We further compared the cortical thickness between preterm neonates and full term neonates, and found the developmental trajectory of cortical thickness in limbic cortex was not impaired by preterm. This implies that the altered hippocampal-limbic covariance may mainly source from the altered hippocampal development. In other word, compared to other neocortical regions, hippocampus exhibited higher degree of vulnerability or plasticity in early extrauterine exposure. In the mnemonic circuitry, which includes a large reciprocal network of regions, hippocampus receives input from almost all neocortical association areas via perirhinal and parahippocampal cortices and finally through the entorhinal cortex (EC) (Van Strien et al., 2009). Thus, the hippocampus has long been considered as a classic example for the study of neuroplasticity as many models of synaptic plasticity have been observed in hippocampal circuits and are thought to be fundamental to learning and memory (Bliss and Schoepfer, 2004; Pastalkova et al., 2006). The high degree of hippocampal plasticity in response to external and internal context alterations, on the other hand, is another saying of high degree of vulnerability (McEwen, 1994). Particularly, when it comes along with some deleterious conditions such as brain injury, impairments, diseases, chronic stress (Bartsch and Wulff, 2015) or the preterm birth which is illustrated in current study. Especially, compared to right hemisphere, winner-take-all analysis indicated that much greater number of left hippocampal vertices established stronger associations with other cortical regions rather than limbic cortex. Most of these vertices, which locate in the lateral hippocampus in left hemisphere, were also the vertices showed faster growth rate in preterm neonates. This suggested that the abnormal structural covariance in left hippocampal hemisphere may also associated with the accelerated growth or catch up process in left hippocampi.

Taken together, we speculate that the asymmetrically accelerated thickness growth exhibited in left hippocampi for preterm neonates may induce two other related asymmetrical developmental trajectories, i.e. the larger left hippocampal volume relative to full term neonates, and the impaired structural covariance between left hippocampi and the limbic cortex. These unexpected asymmetries in preterm neonates, may reveal the molecular and morphological characteristics of neuronal connections according to hippocampal hemisphere (Hutsler and Galuske, 2003), which has proved has further implications for hippocampal function (Shipton et al., 2014; Zaidel, 1995). However, it remains to investigate the specific meaning and the mechanism underlying the asymmetrical developmental trajectory, the persistence of the observed asymmetrical patterns and its long-term consequences in longitudinal studies. This could be an interesting topic for future studies.

### Common or symmetrical impairments induced by preterm birth

Besides the asymmetrical developmental impairments, preterm birth also induced some of the common impairments to hippocampus. Decreased hippocampal thickness were found in both medial and lateral (adjacent to amygdala) parts of the hippocampal head for preterm infants compared to term-born healthy peers in both hemispheres. The hippocampus is known to be critical to learning and memory functions and different portion of the hippocampus in the longitudinal axis functions heterogeneously (Bohbot et al., 2000; Cabeza and Nyberg, 2000; Nadel et al., 2000). The hippocampal head, body, and tail are connected to separate regions of the entorhinal cortex, which conveys processed information from the association cortices to the hippocampus (Hackert et al., 2002). Associations of larger hippocampal volume and better verbal memory performance were found in numerous MRI studies (Jack et al., 1999; Laakso et al., 2000; Tisserand et al., 2000). Specifically, the memory encoding but not retrieval of the verbal memory performance (Lee et al., 2019), may be limited to the head region of the hippocampus. Whereas the recollection of previous experiences, with spatial memory (Maguire et al., 2000) and the degree of psychopathology (Laakso et al., 2001) relies more on the tail part or posterior part of the hippocampus (Kühn and Gallinat, 2014; Lepage et al., 1998; Nadel et al., 2013). Although this hypothesis has been challenged (Schacter and Wagner, 1999), there is evidence that some memory functions in healthy adults (Hackert et al., 2002) are associated primarily with volume deficits in the anterior part of the hippocampus (Hartzell et al., 2016; Langnes et al., 2020). The insufficiently developed hippocampal thickness at hippocampal head in preterm neonates may suggest that the memory encoding process that being supported relatively more by the anterior hippocampus (aHC) was impaired by preterm birth.

### Hippocampal impairments may be induced by other clinical risk factors rather than preterm birth

Of note, the underlying mechanism of accelerated hippocampal growth and hypocovariance with limbic cortex in left hippocampi, and the reduced thickness in hippocampal head for preterm neonates is complex and remains to be clarified, as there have been nearly no previous studies investigating this phenomenon in the preterm cohort. Rather than earlier exposure to extrauterine environment, the impaired hippocampal development may be engendered by other clinical risk factors. One of the main limitations in this study, particularly, is the missing of such important clinical information for the current cohort (not released yet), which raises the concern of group equivalence between full term and preterm neonates. For example, past studies have demonstrated brain injury, including white matter injury, intraventricular hemorrhage (ΓVH) and periventricular leukomalacia or PVL to be an important determinant of hippocampal size from infancy through adolescence (Nosarti et al., 2002; Thompson et al., 2008) For example, Strahle et al. (Strahle et al., 2019) found the injured infants have smaller hippocampi than term and very preterm infants with no or mild brain injuries. Further, past literature proved that (Strahle et al., 2019) very preterm infants with and without brain injury exposed to antenatal steroids had larger hippocampal volumes. Meanwhile, the hippocampus is also known to be sensitive to corticosteroids during its early development (Tombaugh et al., 1992), and exposure to postnatal dexamethasone could impair hippocampal growth (Thompson et al., 2008). Finally, the effects of perinatal clinical risk factors common in preterm infants, including sepsis, necrotizing enterocolitis (NEC), chorioamnionitis and indomethacin exposure, hypotension, chronic lung disease (CLD), and various infections have been extensively studied, with smaller hippocampi and larger ventricles among affected infants (Hatfield et al., 2011).

These findings, although not directly associated with the hippocampal shapes and structural covariance, boost the possibility that the hippocampal growth impairments in premature neonates reported in this study may reflect, at least partially, primary injury or secondary dysmaturational effects of the brain injuries. Future studies will focus on how these clinical risk factors may contribute to the alteration of hippocampal developmental trajectories, particularly the asymmetrical alteration, in preterm neonates.

### Shape analysis comparison

During the third trimester, the human brain undergoes rapid cellular and molecular processes that reshape the structural architecture of cerebral cortex including hippocampus (Ouyang et al., 2019). Shape analysis, as confirmed by prior studies (Joseph et al., 2014; Tae et al., 2011), is an excellent toolbox for representing these regional alterations, and tracking the preterm birth induced abnormal developmental trajectory for hippocampal cortical surfaces. In this study, we performed surface reconstruction via Laplace-Beltrami eigen-projection and properly designed boundary deformations and used thickness measurements during the shape registration for statistical analysis (Shi et al., 2014), which was widely applied in our previous publications (Ge et al., 2015). This method is robust on different input masks of the same structure, and can reconstruct accurate surface representations without introducing artificial oscillations. The final surface is the eigen-projection of the filtered mask boundary that has the correct topology, desired accuracy and smoothness, and was demonstrated to be more sensitive in extracting local information for complex brain regions like hippocampus.

### Other limitations and future directions

The effects of clinical risk factors on hippocampal development is urgently needed in future studies to clarify the compounds effects in the current results. The concept of structural covariance networks describes the interindividual differences in regional brain structure covariation with other brain structures across the population (Alexander-Bloch et al., 2013a). Though the underlying biological meaning of structural covariance is still unclear, it may reflect shared variation in thickness morphology (Alexander-Bloch et al., 2013b) and has been suggested to reflect the synchronized maturational changes medicated by axonal connections forming and reforming over the course of development (Ge et al., 2019). Future studies are needed to compare the structural covariance with other physiologically meaningful connectivity in other modalities such as diffusion tractography and functional connectivity. Shape analysis measures the morphometrical alteration in preterm brain, however little is known about the functional topography of the immature hippocampus (Stoodley and Limperopoulos, 2016). Ongoing studies are needed to better elucidate the impact of these regional alterations in hippocampal development on long-term neurodevelopmental outcomes.

## References

Aggleton, J.P., Pralus, A., Nelson, A.J., Hornberger, M.J.B., 2016. Thalamic pathology and memory loss in early Alzheimer’s disease: moving the focus from the medial temporal lobe to Papez circuit. 139, 1877–1890.

Alexander-Bloch, A., Giedd, J.N., Bullmore, E.J.N.R.N., 2013a. Imaging structural covariance between human brain regions. 14, 322–336.

Alexander-Bloch, A., Raznahan, A., Bullmore, E., Giedd, J.J.J.o.N., 2013b. The convergence of maturational change and structural covariance in human cortical networks. 33, 2889–2899.

Avants, B., Gee, J.C.J.N., 2004. Geodesic estimation for large deformation anatomical shape averaging and interpolation. 23, S139–S150.

Avants, B.B., Tustison, N.J., Song, G., Gee, J.C.J.P.I.C., Laboratory, S., 2009. ANTS: Advanced Open-Source Normalization Tools for Neuroanatomy.

Bajic, D., Moreira, N.C., Wikström, J., Raininko, R.J.A.j.o.n., 2012. Asymmetric development of the hippocampal region is common: a fetal MR imaging study. 33, 513–518.

Ball, G., Boardman, J.P., Rueckert, D., Aljabar, P., Arichi, T., Merchant, N., Gousias, I.S., Edwards, A.D., Counsell, S.J.J.C.c., 2012. The effect of preterm birth on thalamic and cortical development. 22, 1016–1024.

Bartsch, T., Wulff, P., 2015. The hippocampus in aging and disease: from plasticity to vulnerability. Elsevier.

Batalle, D., Hughes, E.J., Zhang, H., Tournier, J.-D., Tusor, N., Aljabar, P., Wali, L., Alexander, D.C., Hajnal, J.V., Nosarti, C.J.N., 2017. Early development of structural networks and the impact of prematurity on brain connectivity. 149, 379–392.

Beauchamp, M.H., Thompson, D.K., Howard, K., Doyle, L.W., Egan, G.F., Inder, T.E., Anderson, P.J.J.B., 2008. Preterm infant hippocampal volumes correlate with later working memory deficits. 131, 2986–2994.

Benjamini, Y., Hochberg, Y.J.J.o.t.R.s.s.s.B., 1995. Controlling the false discovery rate: a practical and powerful approach to multiple testing. 57, 289–300.

Bliss, T., Schoepfer, R.J.S., 2004. Controlling the ups and downs of synaptic strength. 304, 973–974.

Bohbot, V.D., Allen, J.J., Nadel, L.J.A.o.t.N.Y.A.o.S., 2000. Memory deficits characterized by patterns of lesions to the hippocampus and parahippocampal cortex. 911, 355–368.

Cabeza, R., Nyberg, L.J.C.o.i.n., 2000. Neural bases of learning and memory: functional neuroimaging evidence. 13, 415–421.

Cordero Grande, L., Hughes, E.J., Hutter, J., Price, A.N., Hajnal, J.V.J.M.r.i.m., 2018. Three dimensional motion corrected sensitivity encoding reconstruction for multi shot multi slice MRI: application to neonatal brain imaging. 79, 1365–1376.

DuPre, E., Spreng, R.N.J.N.N., 2017. Structural covariance networks across the life span, from 6 to 94 years of age. 1, 302–323.

Duvernoy, H.M., 2005. The human hippocampus: functional anatomy, vascularization and serial sections with MRI. Springer Science & Business Media.

Gahm, J.K., Shi, Y., analysis, A.s.D.N.I.J.M.i., 2018. Riemannian metric optimization on surfaces (RMOS) for intrinsic brain mapping in the Laplace–Beltrami embedding space. 46, 189–201.

Ge, R., Kot, P., Liu, X., Lang, D.J., Wang, J.Z., Honer, W.G., Vila□Rodríguez, F.J.H.b.m., 2019. Parcellation of the human hippocampus based on gray matter volume covariance: Replicable results on healthy young adults. 40, 3738–3752.

Ge, X., Shi, Y., Li, J., Zhang, Z., Lin, X., Zhan, J., Ge, H., Xu, J., Yu, Q., Leng, Y.J.N., 2015. Development of the human fetal hippocampal formation during early second trimester. 119, 33–43.

Gerig, G., Styner, M., Shenton, M.E., Lieberman, J.A., 2001. Shape versus size: Improved understanding of the morphology of brain structures. International Conference on Medical Image Computing and Computer-Assisted Intervention. Springer, pp. 24–32.

Guo, P., Li, Q., Wang, X., Li, X., Wang, S., Xie, Y., Xie, Y., Fu, Z., Zhang, X., Li, S.J.F.i.N., 2020. Structural covariance changes of anterior and posterior hippocampus during musical training in young adults. 14.

Hackert, V., den Heijer, T., Oudkerk, M., Koudstaal, P., Hofman, A., Breteler, M.J.N., 2002. Hippocampal head size associated with verbal memory performance in nondemented elderly. 17, 1365–1372.

Hartzell, J.F., Davis, B., Melcher, D., Miceli, G., Jovicich, J., Nath, T., Singh, N.C., Hasson, U.J.N., 2016. Brains of verbal memory specialists show anatomical differences in language, memory and visual systems. 131, 181–192.

Hatfield, T., Wing, D.A., Buss, C., Head, K., Muftuler, L.T., Davis, E.P.J.A.j.o.o., gynecology, 2011. Magnetic resonance imaging demonstrates long-term changes in brain structure in children born preterm and exposed to chorioamnionitis. 205, 384. e381–384. e388.

Ho, B.-C., Magnotta, V.J.N., 2010. Hippocampal volume deficits and shape deformities in young biological relatives of schizophrenia probands. 49, 3385–3393.

Hutsler, J., Galuske, R.A.J.T.i.n., 2003. Hemispheric asymmetries in cerebral cortical networks. 26, 429–435.

Hwang, H., Rehman, H.Z.U., Lee, S.J.A.S., 2019. 3D U-Net for skull stripping in brain MRI. 9, 569.

Jack, C.R., Petersen, R.C., Xu, Y.C., O’Brien, P.C., Smith, G.E., Ivnik, R.J., Boeve, B.F., Waring, S.C., Tangalos, E.G., Kokmen, E.J.N., 1999. Prediction of AD with MRI-based hippocampal volume in mild cognitive impairment. 52, 1397–1397.

Jacob, F.D., Habas, P.A., Kim, K., Corbett-Detig, J., Xu, D., Studholme, C., Glenn, O.A.J.P.r., 2011. Fetal hippocampal development: analysis by magnetic resonance imaging volumetry. 69, 425–429.

Joseph, J., Warton, C., Jacobson, S.W., Jacobson, J.L., Molteno, C.D., Eicher, A., Marais, P., Phillips, O.R., Narr, K.L., Meintjes, E.M.J.H.b.m., 2014. Three dimensional surface deformation based shape analysis of hippocampus and caudate nucleus in children with fetal alcohol spectrum disorders. 35, 659–672.

Kahn, I., Andrews-Hanna, J.R., Vincent, J.L., Snyder, A.Z., Buckner, R.L.J.J.o.n., 2008. Distinct cortical anatomy linked to subregions of the medial temporal lobe revealed by intrinsic functional connectivity. 100, 129–139.

Khundrakpam, B.S., Lewis, J.D., Reid, A., Karama, S., Zhao, L., Chouinard-Decorte, F., Evans, A.C., Neuroimage, B.D.C.G.J., 2017. Imaging structural covariance in the development of intelligence. 144, 227–240.

Kim, D.J., Park, B., Park, H.J.J.H.b.m., 2013. Functional connectivity based identification of subdivisions of the basal ganglia and thalamus using multilevel independent component analysis of resting state fMRI. 34, 1371–1385.

Kim, S.Y., Liu, M., Hong, S.-J., Toga, A.W., Barkovich, A.J., Xu, D., Kim, H.J.C.C., 2020. Disruption and compensation of sulcation-based covariance networks in neonatal brain growth after perinatal injury. 30, 6238–6253.

Krogsrud, S.K., Tamnes, C.K., Fjell, A.M., Amlien, I., Grydeland, H., Sulutvedt, U., Due Tønnessen, P., Bjørnerud, A., Sølsnes, A.E., Håberg, A.K.J.H.B.M., 2014. Development of hippocampal subfield volumes from 4 to 22 years. 35, 5646–5657.

Kühn, S., Gallinat, J.J.M.p., 2014. Amount of lifetime video gaming is positively associated with entorhinal, hippocampal and occipital volume. 19, 842–847.

Kuklisova-Murgasova, M., Quaghebeur, G., Rutherford, M.A., Hajnal, J.V., Schnabel, J.A.J.M.i.a., 2012. Reconstruction of fetal brain MRI with intensity matching and complete outlier removal. 16, 1550–1564.

Laakso, M.P., Frisoni, G.B., Könönen, M., Mikkonen, M., Beltramello, A., Geroldi, C., Bianchetti, A., Trabucchi, M., Soininen, H., Aronen, H.J.J.B.p., 2000. Hippocampus and entorhinal cortex in frontotemporal dementia and Alzheimer’s disease: a morphometric MRI study. 47, 1056–1063.

Laakso, M.P., Vaurio, O., Koivisto, E., Savolainen, L., Eronen, M., Aronen, H.J., Hakola, P., Repo, E., Soininen, H., Tiihonen, J.J.B.b.r., 2001. Psychopathy and the posterior hippocampus. 118, 187–193.

Langnes, E., Sneve, M.H., Sederevicius, D., Amlien, I.K., Walhovd, K.B., Fjell, A.M.J.H., 2020. Anterior and posterior hippocampus macro and microstructure across the lifespan in relation to memory—A longitudinal study.

Lee, S.-H., Kravitz, D.J., Baker, C.I.J.C.C., 2019. Differential representations of perceived and retrieved visual information in hippocampus and cortex. 29, 4452–4461.

Lepage, M., Habib, R., Tulving, E.J.H., 1998. Hippocampal PET activations of memory encoding and retrieval: the HIPER model. 8, 313–322.

Lerch, J.P., Worsley, K., Shaw, W.P., Greenstein, D.K., Lenroot, R.K., Giedd, J., Evans, A.C.J.N., 2006. Mapping anatomical correlations across cerebral cortex (MACACC) using cortical thickness from MRI. 31, 993–1003.

Li, X., Pu, F., Fan, Y., Niu, H., Li, S., Li, D.J.F.i.h.n., 2013. Age-related changes in brain structural covariance networks. 7, 98.

Liu, M., Lepage, C., Jeon, S., Flynn, T., Yuan, S., Kim, J., Toga, A.W., Barkovich, A.J., Xu, D., Evans, A.C., 2019. A Skeleton and Deformation Based Model for Neonatal Pial Surface Reconstruction in Preterm Newborns. 2019 IEEE 16th International Symposium on Biomedical Imaging (ISBI 2019). IEEE, pp. 352–355.

Maguire, E.A., Mummery, C.J., Büchel, C.J.H., 2000. Patterns of hippocampal cortical interaction dissociate temporal lobe memory subsystems. 10, 475–482.

Makropoulos, A., Robinson, E.C., Schuh, A., Wright, R., Fitzgibbon, S., Bozek, J., Counsell, S.J., Steinweg, J., Vecchiato, K., Passerat-Palmbach, J.J.N., 2018. The developing human connectome project: A minimal processing pipeline for neonatal cortical surface reconstruction. 173, 88–112.

Maller, J.J., Welton, T., Middione, M., Callaghan, F.M., Rosenfeld, J.V., Grieve, S.M.J.S.r., 2019. Revealing the hippocampal connectome through super-resolution 1150-direction diffusion MRI. 9, 1–13.

McEwen, B.S., 1994. The plasticity of the hippocampus is the reason for its vulnerability. Seminars in Neuroscience. Elsevier, pp. 239–246.

Mechelli, A., Friston, K.J., Frackowiak, R.S., Price, C.J.J.J.o.N., 2005. Structural covariance in the human cortex. 25, 8303–8310.

Mills, K.L., Bathula, D., Costa Dias, T.G., Iyer, S.P., Fenesy, M.C., Musser, E.D., Stevens, C.A., Thurlow, B.L., Carpenter, S.D., Nagel, B.J.J.F.i.p., 2012. Altered cortico-striatal–thalamic connectivity in relation to spatial working memory capacity in children with ADHD. 3, 2.

Nadel, L., Hoscheidt, S., Ryan, L.R.J.J.o.c.n., 2013. Spatial cognition and the hippocampus: the anterior–posterior axis. 25, 22–28.

Nadel, L., Samsonovich, A., Ryan, L., Moscovitch, M.J.H., 2000. Multiple trace theory of human memory: computational, neuroimaging, and neuropsychological results. 10, 352–368.

Nordin, K., Persson, J., Stening, E., Herlitz, A., Larsson, E.M., Söderlund, H.J.H., 2018. Structural whole brain covariance of the anterior and posterior hippocampus: Associations with age and memory. 28, 151–163.

Nosarti, C., Al□Asady, M.H., Frangou, S., Stewart, A.L., Rifkin, L., Murray, R.M.J.B., 2002. Adolescents who were born very preterm have decreased brain volumes. 125, 1616–1623.

Nosarti, C., Froudist□Walsh, S.J.D.M., Neurology, C., 2016. Alterations in development of hippocampal and cortical memory mechanisms following very preterm birth. 58, 35–45.

Nosarti, C., Nam, K.W., Walshe, M., Murray, R.M., Cuddy, M., Rifkin, L., Allin, M.P.J.N.C., 2014. Preterm birth and structural brain alterations in early adulthood. 6, 180–191.

Ouyang, M., Jeon, T., Sotiras, A., Peng, Q., Mishra, V., Halovanic, C., Chen, M., Chalak, L., Rollins, N., Roberts, T.P.J.P.o.t.N.A.o.S., 2019. Differential cortical microstructural maturation in the preterm human brain with diffusion kurtosis and tensor imaging. 116, 4681–4688.

Pastalkova, E., Serrano, P., Pinkhasova, D., Wallace, E., Fenton, A.A., Sacktor, T.C.J.s., 2006. Storage of spatial information by the maintenance mechanism of LTP. 313, 1141–1144.

Pedraza, O., Bowers, D., Gilmore, R.J.J.o.t.I.N.S., 2004. Asymmetry of the hippocampus and amygdala in MRI volumetric measurements of normal adults. 10, 664–678.

Perlman, J.M.J.P., 2001. Neurobehavioral deficits in premature graduates of intensive care—potential medical and neonatal environmental risk factors. 108, 1339–1348.

Persson, J., Spreng, R.N., Turner, G., Herlitz, A., Morell, A., Stening, E., Wahlund, L.-O., Wikström, J., Söderlund, H.J.N., 2014. Sex differences in volume and structural covariance of the anterior and posterior hippocampus. 99, 215–225.

Poppenk, J., Evensmoen, H.R., Moscovitch, M., Nadel, L.J.T.i.c.s., 2013. Long-axis specialization of the human hippocampus. 17, 230–240.

Raaijmakers, A., Allegaert, K., 2016. Catch-up growth in former preterm neonates: no time to waste. Multidisciplinary Digital Publishing Institute.

Raznahan, A., Lerch, J.P., Lee, N., Greenstein, D., Wallace, G.L., Stockman, M., Clasen, L., Shaw, P.W., Giedd, J.N.J.N., 2011. Patterns of coordinated anatomical change in human cortical development: a longitudinal neuroimaging study of maturational coupling. 72, 873–884.

Ronneberger, O., Fischer, P., Brox, T., 2015. U-net: Convolutional networks for biomedical image segmentation. International Conference on Medical image computing and computer-assisted intervention. Springer, pp. 234–241.

Sakaguchi, Y., Sakurai, Y.J.B.B.R., 2017. Left–right functional asymmetry of ventral hippocampus depends on aversiveness of situations. 325, 25–33.

Schacter, D.L., Wagner, A.D.J.H., 1999. Medial temporal lobe activations in fMRI and PET studies of episodic encoding and retrieval. 9, 7–24.

Schmidt-Kastner, R., Freund, T.J.N., 1991. Selective vulnerability of the hippocampus in brain ischemia. 40, 599–636.

Shah, A., Jhawar, S.S., Goel, A.J.J.o.C.N., 2012. Analysis of the anatomy of the Papez circuit and adjoining limbic system by fiber dissection techniques. 19, 289–298.

Shi, Y., Lai, R., Morra, J.H., Dinov, I., Thompson, P.M., Toga, A.W.J.I.t.o.m.i., 2010. Robust surface reconstruction via Laplace-Beltrami eigen-projection and boundary deformation. 29, 2009–2022.

Shi, Y., Lai, R., Wang, D.J., Pelletier, D., Mohr, D., Sicotte, N., Toga, A.W.J.I.t.o.m.i., 2014. Metric optimization for surface analysis in the Laplace-Beltrami embedding space. 33, 1447–1463.

Shinohara, M.L., Kim, H.-J., Kim, J.-H., Garcia, V.A., Cantor, H.J.P.o.t.N.A.o.S., 2008. Alternative translation of osteopontin generates intracellular and secreted isoforms that mediate distinct biological activities in dendritic cells. 105, 7235–7239.

Shipton, O.A., El-Gaby, M., Apergis-Schoute, J., Deisseroth, K., Bannerman, D.M., Paulsen, O., Kohl, M.M.J.P.o.t.N.A.o.S., 2014. Left–right dissociation of hippocampal memory processes in mice. 111, 15238–15243.

Smitthimedhin, A., Whitehead, M.T., Bigdeli, M., Nino, G., Perez, G., Otero, H.J.J.C.i., 2018. MRI determination of volumes for the upper airway and pharyngeal lymphoid tissue in preterm and term infants. 50, 51–56.

Stoodley, C.J., Limperopoulos, C., 2016. Structure–function relationships in the developing cerebellum: evidence from early-life cerebellar injury and neurodevelopmental disorders. Seminars in Fetal and Neonatal Medicine. Elsevier, pp. 356–364.

Strahle, J.M., Triplett, R.L., Alexopoulos, D., Smyser, T.A., Rogers, C.E., Limbrick Jr, D.D., Smyser, C.D.J.N.C., 2019. Impaired hippocampal development and outcomes in very preterm infants with perinatal brain injury. 22, 101787.

Tae, W., Kim, S., Lee, K., Nam, E., Choi, J., Park, J.J.A.j.o.n., 2011. Hippocampal shape deformation in female patients with unremitting major depressive disorder. 32, 671–676.

Tanaka, K.Z.J.N.R., 2020. Heterogeneous representations in the hippocampus.

Thompson, D.K., Adamson, C., Roberts, G., Faggian, N., Wood, S.J., Warfield, S.K., Doyle, L.W., Anderson, P.J., Egan, G.F., Inder, T.E.J.N., 2013. Hippocampal shape variations at term equivalent age in very preterm infants compared with term controls: perinatal predictors and functional significance at age 7. 70, 278–287.

Thompson, D.K., Loh, W.Y., Connelly, A., Cheong, J.L., Spittle, A.J., Chen, J., Kelly, C.E., Inder, T.E., Doyle, L.W., Anderson, P.J.J.P.r., 2020. Basal ganglia and thalamic tract connectivity in very preterm and full-term children; associations with 7-year neurodevelopment. 87, 48–56.

Thompson, D.K., Wood, S.J., Doyle, L.W., Warfield, S.K., Egan, G.F., Inder, T.E.J.H., 2009. MR□determined hippocampal asymmetry in full□term and preterm neonates. 19, 118–123.

Thompson, D.K., Wood, S.J., Doyle, L.W., Warfield, S.K., Lodygensky, G.A., Anderson, P.J., Egan, G.F., Inder, T.E.J.A.o.n., 2008. Neonate hippocampal volumes: prematurity, perinatal predictors, and 2 year outcome. 63, 642–651.

Tisserand, D., Visser, P., Van Boxtel, M., Jolles, J.J.N.o.a., 2000. The relation between global and limbic brain volumes on MRI and cognitive performance in healthy individuals across the age range. 21, 569–576.

Tombaugh, G.C., Yang, S.H., Swanson, R.A., Sapolsky, R.M.J.J.o.n., 1992. Glucocorticoids exacerbate hypoxic and hypoglycemic hippocampal injury in vitro: biochemical correlates and a role for astrocytes. 59, 137–146.

Uematsu, A., Matsui, M., Tanaka, C., Takahashi, T., Noguchi, K., Suzuki, M., Nishijo, H.J.P.o., 2012. Developmental trajectories of amygdala and hippocampus from infancy to early adulthood in healthy individuals. 7, e46970.

Van Strien, N., Cappaert, N., Witter, M.J.N.r.n., 2009. The anatomy of memory: an interactive overview of the parahippocampal–hippocampal network. 10, 272–282.

Vann, S.D.J.E., 2013. Dismantling the Papez circuit for memory in rats. 2, e00736.

Vasung, L., Lepage, C., Radoš, M., Pletikos, M., Goldman, J.S., Richiardi, J., Raguž, M., Fischi-Gómez, E., Karama, S., Huppi, P.S.J.F.i.n., 2016. Quantitative and qualitative analysis of transient fetal compartments during prenatal human brain development. 10, 11.

Wang, Z., Zou, N., Shen, D., Ji, S., 2020. Non-Local U-Nets for Biomedical Image Segmentation. AAAI, pp. 6315–6322.

Wolf, H., Grunwald, M., Kruggel, F., Riedel-Heller, S., Angerhöfer, S., Hojjatoleslami, A., Hensel, A., Arendt, T., Gertz, H.-J.J.N.o.a., 2001. Hippocampal volume discriminates between normal cognition; questionable and mild dementia in the elderly. 22, 177–186.

Wright, I., McGuire, P., Poline, J.-B., Travere, J., Murray, R., Frith, C., Frackowiak, R., Friston, K.J.N., 1995. A voxel-based method for the statistical analysis of gray and white matter density applied to schizophrenia. 2, 244–252.

Wu, D., Chang, L., Akazawa, K., Oishi, K., Skranes, J., Ernst, T., Oishi, K.J.D.i.b., 2017. Change-point analysis data of neonatal diffusion tensor MRI in preterm and term-born infants. 12, 453–458.

Yushkevich, P.A., Gao, Y., Gerig, G., 2016. ITK-SNAP: An interactive tool for semiautomatic segmentation of multi-modality biomedical images. 2016 38th Annual International Conference of the IEEE Engineering in Medicine and Biology Society (EMBC). IEEE, pp. 3342–3345.

Zaidel, D.W.J.B.r., 1995. The case for a relationship between human memory, hippocampus and corpus callosum. 28, 51–57.

Zhang, D., Snyder, A.Z., Fox, M.D., Sansbury, M.W., Shimony, J.S., Raichle, M.E.J.J.o.n., 2008. Intrinsic functional relations between human cerebral cortex and thalamus. 100, 1740–1748.

